# Characterization of the alfalfa pollen virome

**DOI:** 10.64898/2026.02.17.706418

**Authors:** Lev G. Nemchinov, Sam Grinstead, Olga A. Postnikova, Brian M. Irish

**Author notes:** Corresponding author: Nemchinov LG.

## Abstract

Vertical transmission of plant pathogenic viruses is an important component of viral persistence, survival, and spread in agricultural production systems. This type of transmission is of considerable economic significance as it can cause major crop losses by serving as the initial focus of infection for future epidemics. Vertical transmission occurs when a virus is passed on to offspring either by direct invasion of the developing seed embryo from infected mother plants or through infected pollen grains after fertilization. We have recently demonstrated by high throughput sequencing that mature seeds of the agriculturally important forage crop alfalfa (*Medicago sativa* L.) are associated with a broad range of viruses some of which could potentially spread over long distances via seed. Aside from alfalfa mosaic virus, little is currently known about viral transmission via alfalfa pollen and its role in the epidemiology in this crop. This research was conducted to screen the pollen obtained from unique alfalfa genotypes for the presence of pathogenic viruses and their potential for dissemination. The plants from which the pollen was collected were alfalfa genotypes selected for fungal plant disease resistance and agronomic performance in a USDA ARS pre-breeding program in Prosser, WA.

## Introduction

Pollen transmission of plant viruses is a part of the vertical transmission from parents to offspring. It can occur when virus in the pollen grains infects the ovule during fertilization, which leads to infected embryo, seeds, and eventually to the virus-infected seedlings and mature plants [1, 2, 3, 4]. Although majority of pathogenic viruses are excluded from apical meristems, vertical transmission to the host progeny could also take place via virus invasion of the meristematic tissues and subsequently of all plant reproductive organs [5, 6]. As pollen is known to travel long distances via airborne particles or by insect pollinators [7, 8], the transmission of pathogenic viruses can result in expansive dissemination and their introduction into the new areas.

The dissemination of viruses in pollen is common and has been reported in several agricultural crops [9,10]. However, the current experimental knowledge on pollen transmission of plant viruses in the agriculturally important alfalfa (*Medicago sativa* L.) forage crop is limited to alfalfa mosaic virus (AMV), [11,12]. In greenhouse experiments, Frosheiser [11] showed that the frequency of AMV transmission to alfalfa seeds through pollen ranged from 0.5 to 26.5%.

The first virome of alfalfa seeds recently obtained by high throughput sequencing (HTS) indicated a presence of sequencing reads belonging to 27 viruses from 10 different families [13]. The prevailing species were AMV, ubiquitous partiti, amalga and similarly persistent snake river alfalfa virus (SRAV). Several other viruses not previously associated with alfalfa seeds were also identified [13].

At the moment, the nature and mechanism of alfalfa seed contamination by these viruses is unknown. While they could enter into the seed parts of maternal origin indirectly, through infection of reproductive organs before embryogenesis [2], viral access could also occur via direct invasion of the embryo during embryogenesis, for instance by way of infected pollen [1, 2, 3, 4]. This study highlights the significant role that pollen infection may play in the vertical transmission of plant viruses in alfalfa.

## Methods

### Plant material

Pollen was collected from 15 different alfalfa genotypes previously selected in an in-house pre-breeding program for disease resistance and good agronomic performance (Additional file 1). Plants were carried over from the 2024 growing season specifically for pollen collection from previous year polycrosses. Original plant germplasm came from the USDA ARS National Plant Germplasm System alfalfa collection. Once regrowth began in the early summer alfalfa plants were covered by a 6.1×6.1×1.8 m insect-proof cages for exclusion of potential pollinators. Field site was on Washington State University Irrigated Agriculture Research and Extension Center where the USDA-ARS Plant Germplasm Introduction and Testing Research Unit (PGITRU) has a worksite. Branches with racemes and many recently opened flowers were collected from four clonally propagated plants of each of the 15 genotypes. These flower ‘bouquets’ were then transferred to the laboratory, kept turgid with base in water and used to collect pollen into 1.5 ml microfuge tubes. On average, approximately 750 flowers per genotype were used for pollen collection. Individual flowers were tripped with a 1 ml pipette tip cut at an angle (Additional File 2). The methodology yielded pure pollen samples, free of visible contamination from vegetative tissues (Fig. 1).

**Figure 1.**
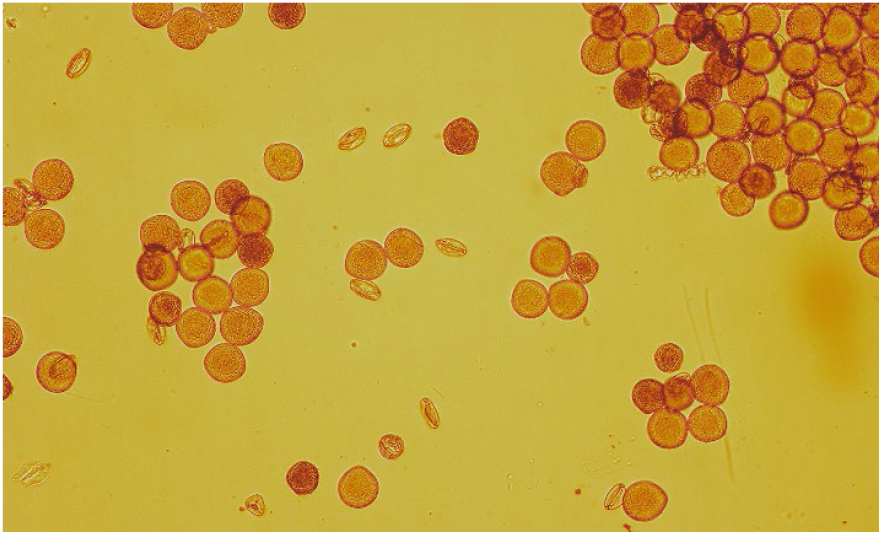
Alfalfa pollen samples obtained by manually tripping individual flowers, without visible contamination from vegetative tissues (Leica DM2000, 40X).

At regular intervals (e.g., every 50-100 flowers) pollen at the end of the pipette tip was transferred with a pipettor to 1ml of Trizol Reagent (Thermo Fisher Scientific Inc., Waltham, MA, USA) by aspirating and expelling liquid a few times and briefly stored at 4°C before RNA extraction (Additional File 3). Several methods were tested for pollen storage and preservation after collection and prior to the RNA extraction, including the use of RNAlater and TRIzol reagents, both supplied by ThermoFisher Scientific (Waltham, MA, USA). Microscopic observations showed that in RNAlater pollen grains looked wrinkled and shrunken after a day of storage, while pollen samples in TRIzol Reagent looked for the mostly intact and free of any visible contaminants (Fig. 1; Additional File 4).

### Total RNA extraction, RNA sequencing and RT-PCR

Before RNA extraction, pollen samples stored in TRIzol Reagent were disrupted using FastPrep-24 5G homogenizer (MP Biomedicals, Irvine, CA, USA) to access viruses inside the pollen grains. Pollen was transferred to 2 ml FastPrep tubes containing lysing matrix D (MP Biomedicals, Irvine, CA, USA) and homogenized 5 times at maximum speed (10m/sec) for 1 minute each time. Once the samples were examined under the microscope and found to be sufficiently lysed and disrupted with their cytoplasm released (Additional File 5), they were processed for total RNA extraction, following the manufacturer’s protocol for TRIzol Reagent. After completion, the final eluate was additionally processed through Qiagen’s RNeasy Plant Mini Kit for RNA Extraction (Qiagen Inc., Germantown, MD USA). RNA libraries were prepared using the ribosomal RNA depletion method and were sequenced on an Illumina NovaSeq X platform (PE150) by CD Genomics (Shirley, NY, USA). All 15 samples were sequenced in a dedicated single lane to avoid potential cross-contamination from different projects. The RNA sequencing depth was 6Gb per sample. Reverse transcription–polymerase chain reactions (RT-PCR) were performed using the Titan One Tube RT-PCR System according to the manufacturer’s directions (Roche Diagnostics, Mannheim, Germany). Primers specific to each tested virus were designed based on the results of the HTS and are shown in the Additional File 6. To ensure a complete absence of viral contamination (if used as controls, “healthy” alfalfa tissues may contain a variety of viral infections [14]), a sterile RNAse-free water was used in control RT-PCR reactions along with additional no RT reaction controls as recommended by the manufacturer. The RT-PCRs were carried out in two technical replications.

### Bioinformatic analysis

Bioinformatics analyses were performed as previously described [13, 15]. Briefly, sequence reads were trimmed using Trimmomatic [16] then assembled with SPAdes [17]. The resulting contigs were screened using BLASTx searches [18] against a virus database containing all plant virus protein sequences from the NCBI RefSeq database (https://www.ncbi.nlm.nih.gov/refseq/). The resulting potential plant viral hits were searched once again using BLASTx against the full NCBI nr protein database. BBMap [19] was used to generate sequencing coverage values for the final hits. Insect contamination in the pollen was estimated with Kraken2 [20] using the core-nt database to classify the trimmed HTS read pairs (https://benlangmead.github.io/aws-indexes/k2). Samples averaged 93.9% classified reads.

## Results and Discussion

In total, sequencing reads belonging to 22 viruses were found across all pollen samples from 15 different genotypes (Fig. 2, Table 1, Additional file 7).

**Table 1.** A list of viruses identified by HTS in alfalfa pollen collected from 15 different germplasm sources.

| Virus names |  | Proposed taxonomy | *№ alfalfa genotypes |
| --- | --- | --- | --- |
| 1. | Alfalfa cytorhabdovirus 1 | *Rhabdoviridae, Cytorhabdovirus* | 7 |
| 2. | Alfalfa cytorhabdovirus 2 | *Rhabdoviridae, Cytorhabdovirus* | 1 |
| 3. | Alfalfa deltapartitivirus | *Partitiviridae, Deltapartitivirus* | 3 |
| 4. | Alfalfa nucleorhabdovirus 1 | *Rhabdoviridae, Betanucleorhabdovirus* | 12 |
| 5. | Alfalfa toti-like virus 1 | *Orthototiviridae, unclassified* | 1 |
| 6. | Alfalfa-associated picorna-like virus 2 | *Picornavirales, Iflaviridae, Iflavirus* | 4 |
| 7. | Alfamovirus AMV | *Bromoviridae, Alfamovirus* | 15 |
| 8. | Allexivirus sigmamedicagonis (alfalfa virus S) | *Alphaflexiviridae, Allexivirus* | 4 |
| 9. | Bean leafroll virus | *Tombusviridae, Luteovirus* | 1 |
| 10. | Beet cryptic virus 1 | *Partitiviridae, Alphacryptovirus* | 1 |
| 11. | Big Cypress virus | *Orbivirus* | 1 |
| 12. | Caulimovirus MSa3 | *Caulimoviridae, unclassified* | 5 |
| 13. | Caulimovirus spp. | *Caulimoviridae, Caulimovirus* | 15 |
| 14. | Medicago sativa alphapartitivirus 1 | *Partitiviridae, Alphapartitivirus* | 5 |
| 15. | Medicago sativa alphapartitivirus 2 | *Partitiviridae, Alphapartitivirus* | 5 |
| 16. | Medicago sativa amalgavirus 1 | *Amalgaviridae, Unclassified* | 5 |
| 17. | Medicago sativa deltapartitivirus 1 | *Partitiviridae, Deltapartitivirus* | 2 |
| 18. | Pea streak virus | *Betaflexiviridae, Carlavirus* | 15 |
| 19. | Pepper-associated picorna-like virus | *Picornavirales, unclassified* | 8 |
| 20. | Red clover vein mosaic virus | *Betaflexiviridae, Carlavirus* | 2 |
| 21. | Snake River alfalfa virus | *unclassified* | 3 |
| 22. | Alfalfa-associated nucleorhabdovirus | *Rhabdoviridae, Betanucleorhabdovirus* | 1 |

\*The number of alfalfa genotypes in which individual viruses were identified.

**Figure 2.**
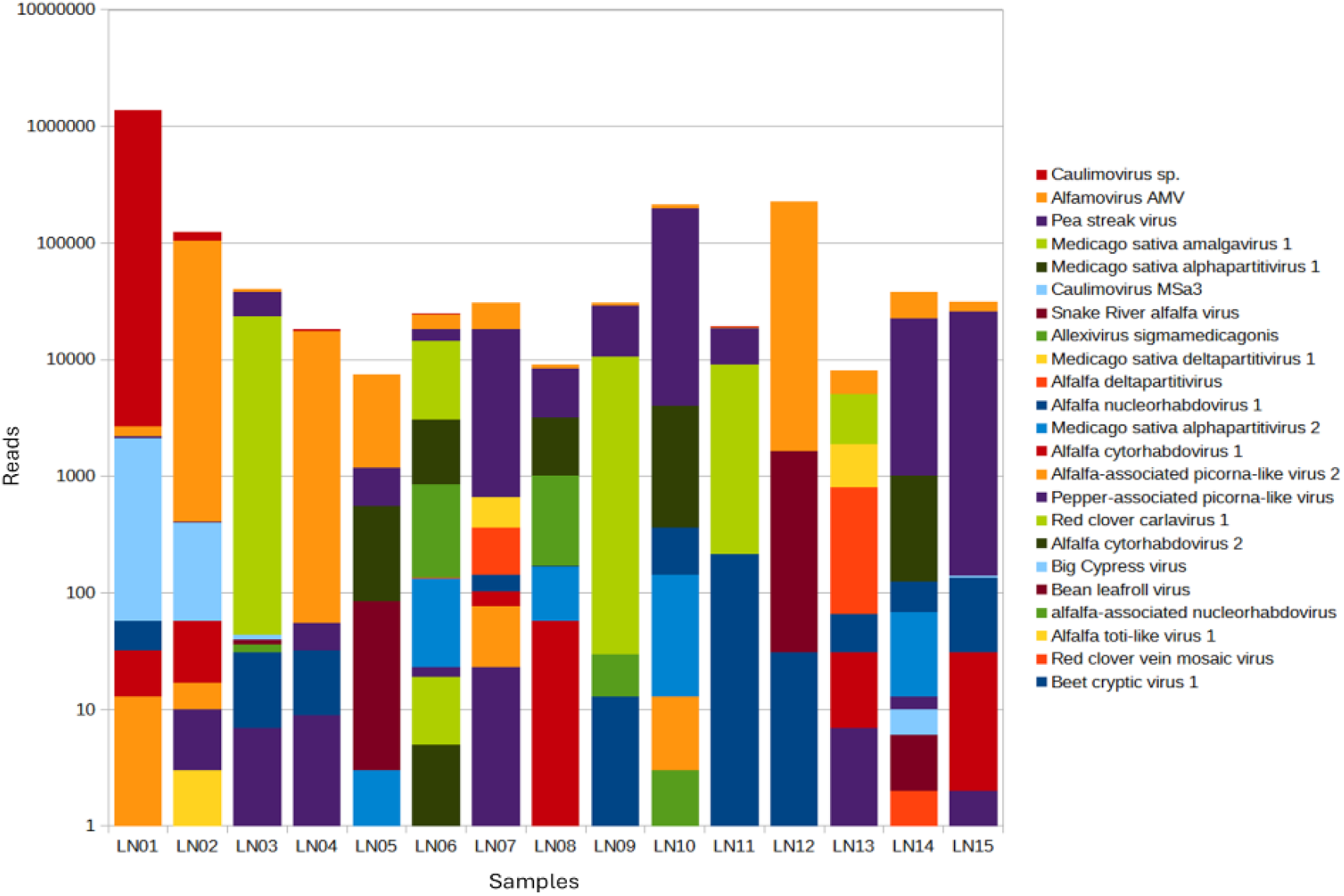
Distribution of viral communities among alfalfa pollen samples collected from 15 different genotypes.

Out of this number, 14 were previously detected in alfalfa seeds [13]. Predictably, these viruses included AMV, partiti, and -amalga viruses, SRAV, and alfalfa virus S (AVS). While AMV is known to be pollen and seed transmitted, so far there has been no experimental evidence of alfalfa pollen contamination by members of the families *Partitiviridae, Amalgoviridae, Alphaflexiviridae* (AVS), and by taxonomically unclear SRAV, even though pollen infection could be assumed based on the seed transmissibility of these viruses [13, 21].

More compelling is that reads of bean leafroll virus (BLRV) and pea streak virus (PeSV), which traditionally are not considered seedborne viruses, were identified in one and in all 15 samples, respectively (Table 1, Additional file 7). This is the third time we were able to detect these two viruses in alfalfa reproductive germplasm [13, 21]. While both viruses are considered of minor importance in alfalfa, they are aphid transmitted and serious pathogens on other legume crops [22]. Consistent detection of cyto- and – nucleorhabdoviral sequencing reads in alfalfa seeds, vegetative tissues and now in pollen samples, may indicate that they potentially represent endogenous viral elements (EVEs) integrated into the plant’s genome [23]. The integrated nucleocapsid protein genes of cytorhabdoviruses were previously found in the genomes of 9 plant families although the mechanism by which the viral RNA sequences were converted to DNA and incorporated into plant genomes remains unknown [24, 25]. Hypothetically, the integration could be facilitated by reverse transcriptase encoded by other elements, such as pararetroviruses or retrotransposons [24]. Pararetroviruses of *Caulimoviridae* family, *Caulimovirus* spp. were identified in all 15 samples (Table 1). The EVEs of two dsDNA reverse-transcribing virus plant pararetroviruses of this family were previously found to be stable constituents of the alfalfa genome [23].

Equally interesting is identification in alfalfa pollen samples of red clover vein mosaic virus (RCVMV), a member of the genus *Carlavirus*, family *Betaflexiviridae*. While RCVMV is known to infect alfalfa [26, 27], to our knowledge, its vertical transmission in the crop has not been reported. In alfalfa, RCVMV can cause reduced forage biomass and quality through symptoms like crinkling, interveinal mosaic, small leaves, yellowing, stunting, and mottling [26]. In alfalfa breeding programs for developing winter hardiness, RCVMV caused premature death of some breeding lines [22]. Alfalfa can also be a reservoir for RCVMV leading to the spread of the virus to other important legume crops that are also susceptible to this pathogen [22].

One of the pollen samples contained sequencing reads of beet cryptic virus (BCV), the first discovered member of the family *Partitiviridae* to infect alfalfa [28]. Biological effects of BCV on alfalfa are unclear, although in sugar beet BCV infection was associated with significantly reduced root and sugar yields [29].

Sequencing reads with high-scoring match (>99% nucleotide identity) to the pepper-associated picorna-like virus (PAPLV) were identified in 8 alfalfa pollen samples. Except for the nucleotide sequence (PP728253.1), we could not find any information on the biology or phylogeny of PAPLV. Although pepper plants in Korea are infected by a variety of viruses including economically important picorna-like viruses [30], PAPLV is not listed among them.

Several other viruses found in pollen could have originated from contamination by arthropods (big Cypress virus, alfalfa-associated picorna-like virus 2), and/or from fungal or protozoan hosts (alfalfa toti-like virus 1). In this regard, it is worth noting that the method used for collecting pollen samples prevented contamination from vegetative tissues (see Methods) and the only potential contaminants could derive from insects feeding on alfalfa flowers, or by fungal hosts or protozoans carried by pollinators.

Although aphids are the primary insect vectors of AMV, BLRV, PeSV, and RCVMV, all of which were found in pollen samples, they do not feed on pollen and mostly congregate on stems and leaves. This was reflected in the very few number of reads associated with pea aphid (*Acyrthosiphon pisum*), one of the most efficient vectors of AMV, BLRV, PeSV and RCVMV (on average, 1.34874E-06 of the total reads per sample; Additional file 8). Yet, it cannot be completely ruled out that BLRV, PeSV, and RCVMV could potentially originate from aphid debris and/or body parts. It is especially true of BLRV and RCVMV, which were associated with a limited number of samples and sequencing reads. The same could possibly apply to SRAV as the virus was previously reported in Western flower thrips (*Frankliniella occidentalis*) feeding on alfalfa stands [31], and limited number of *F. occidentalis* reads were identified in the pollen samples (on average, 0.0001 of the total reads per sample; Additional file 8). However, given that our previous results established the presence of SRAV in surface-sterilized alfalfa seeds and seedlings germinated from them, we believe that the detection of the virus in the pollen grains is authentic.

On the contrary, PeSV reads were identified in each of the 15 tested samples, which is likely to be related to the viral presence on the surface or interior of the pollen grains. However, whether the viruses were in insect debris, a small amount of which could pollute the pollen samples, or in/on pollen grains themselves, is likely irrelevant because virus-carrying insects contaminating pollen samples could easily be dispersed concomitantly thus causing viruses to spread throughout the crop.

We used reverse transcription–polymerase chain reactions to confirm the presence of several arbitrarily chosen viruses that were initially identified by sequencing in alfalfa pollen samples (Fig.2). The RT-PCR assays were conducted on samples identified as positive for each respective virus via HTS. Primers for RT-PCRs were designed based on the obtained HTS contigs (Additional file 6). The viruses included AVS, BLRV, PeSV, SRAV, PAPLV, BCV, RCVMV, ANRV 1, Medicago sativa amalgavirus 1 (MsAV1), and Medicago sativa alphapartitivirus 1 (MsAPV1). The RT-PCR products were successfully amplified from samples that tested positive for every virus via HTS, excluding PAPLV, RCVMV, and BCV (Fig.3).

**Figure 3.**
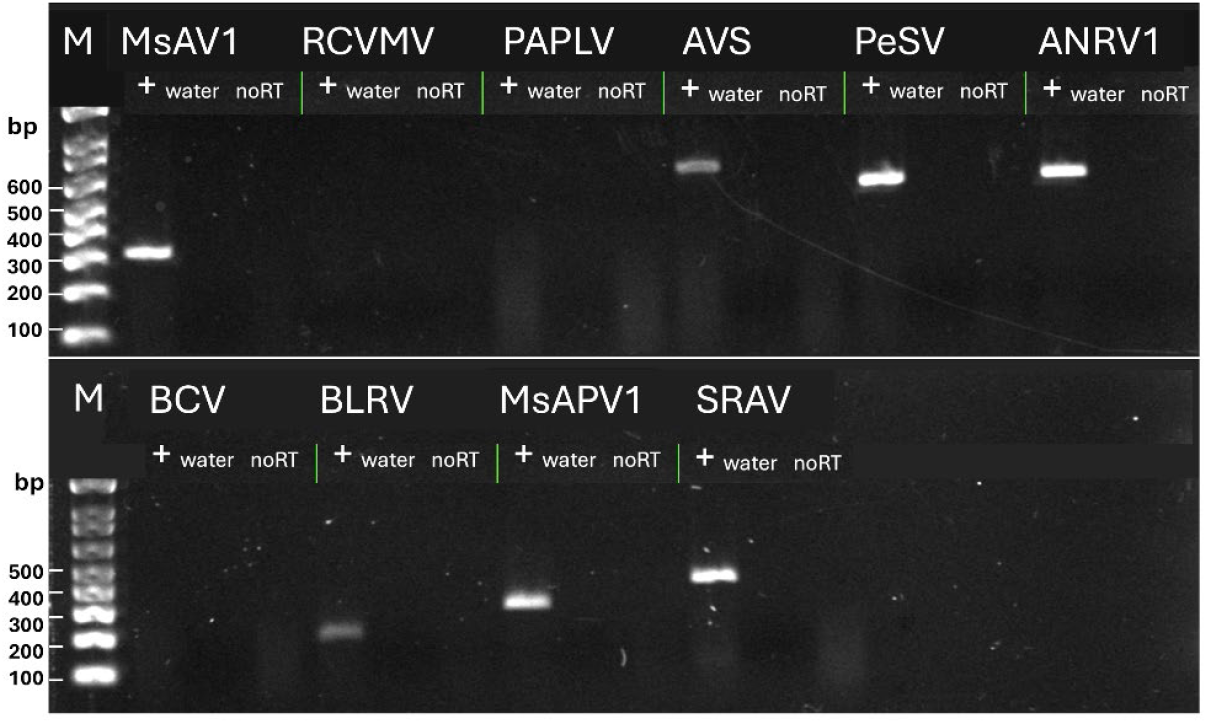
Reverse Transcription Polymerase Chain Reaction (RT-PCR) to validate the presence of arbitrarily selected viral sequences in alfalfa pollen samples. “+” indicates RT-PCR product obtained from a pollen sample; “water” indicates a control reaction with sterile RNAse-free water added instead of a pollen sample; “no RT” indicates a control reaction with all components present except the RT enzyme mix. M, 1kb DNA ladder, (Thermo Fisher Scientific Inc., Waltham, MA USA); MsAPV, Medicago sativa alphapartitivirus; RCVMV, red clover vein mosaic virus; PAPLV, pepper-associated picorna-like virus; AVS, alfalfa virus S; PeSV, pea streak virus; ANRV, alfalfa nucleorhabdovirus; BCV, beet cryptic virus; BLRV, bean leafroll virus; MsAPV, Medicago sativa alphapartitivirus; SRAV, Snake River alfalfa virus.

It is possible that the quantity of viral genetic material for the latter three viruses in the samples was below the threshold for reliable detection via RT-PCR, given the limited number of sequencing reads observed.

## Conclusion

Based on our current information, this is the first study of the alfalfa pollen virome carried out by HTS technology. As of today, the experimental knowledge on alfalfa pollen infection with plant viruses was limited to AMV [11, 12]. In this study, in addition to AMV, we identified sequencing reads belonging to several other viruses infecting and/or contaminating alfalfa pollen grains, including partiti- and -amalga viruses, SRAV, AVS, BLRV, PeSV, RCVMV, PAPLV, and other species. The presence of the arbitrarily chosen viruses (SRAV, AVS, BLRV, PeSV, ANRV, MsAPV1, and MsAV1) in alfalfa pollen samples was also confirmed by RT-PCR with virus-specific primers. These results are consistent with the original findings on alfalfa seed virome obtained by high throughput sequencing [13] and may point to pollen infection as a potential cause of vertical virus transmission.

Vertical transmission introduces a virus into a population at an early age, which can lead to its wider dissemination and subsequent epidemics [2, 32]. It is also important to point out that alfalfa can be a reservoir host for viruses, which can result in substantial losses for other crops [22]. Finally, since virus infections are known to negatively affect reproductive characteristics of the pollen grains and their overall yield [33, 34, 35], it is anticipated that presence of viruses in pollen may further reduce its performance thus possibly impacting seed production.

The practical significance of this study also relates to the specific tested genotypes, which were originally sourced from the USDA-ARS National Plant Germplasm System (NPGS). The genotypes were selected and are being used in pre-breeding efforts to improve alfalfa for fungal disease resistance and good agronomic performance. Finding viral plant pathogens associated with the pollen, including some with reported vertical transmission, is important information to consider as improved selections progress.

## List of abbreviations

AVS: alfalfa virus S
ACRV: alfalfa cytorhabdovirus
ANRV: alfalfa nucleorhabdovirus
BCV: beet cryptic virus
BLRV: bean leafroll virus
HTS: High-throughput sequencing
MsAV: Medicago sativa amalgavirus
MsAPV: Medicago sativa alphapartitivirus
NPGS: National Plant Germplasm System
PAPLV: pepper-associated picorna-like virus
PeSV: pea streak virus
RCVMV: red clover vein mosaic virus
SRAV: Snake River alfalfa virus
RT-PCR: reverse transcription-polymerase chain reaction

## Declarations

### Contribution

LGN: concept, data analysis, wet lab, and first draft of the manuscript; BMI: concept, genotype selection, samples collection and evaluation. SG: bioinformatics and data analysis; OAP: wet lab, bioinformatics, and data analysis. All authors contributed to the editing of the final version of the manuscript and approved it for publication.

### Data Availability Statement

All sequences mentioned in the text are included in the supplementary materials.

The raw metadata were also submitted to the NCBI’s GenBank (accession number pending).

### Ethics approval

Not applicable

### Consent for publication

All authors consent to the publication of the manuscript

## Additional files

**Additional file 1**: Pollen was collected from 15 different alfalfa genotypes maintained in the research facility of Plant Germplasm Introduction and Testing Research Unit, in Prosser, WA.

**Additional file 2:** Individual flowers were tripped with 1 ml pipette tip cut at an angle for pollen collection.

**Additional file 3**. Individual pollen samples were stored in Trizol Reagent (Thermo Fisher Scientific Inc., Waltham, MA USA).

**Additional file 4**. Comparison between pollen samples stored in RNAlater (A) and TRIzol reagents (B) (both supplied by Thermo Fisher Scientific (Waltham, MA USA). Olympus BX53; 40X.

**Additional file 5**. Alfalfa pollen samples disrupted in FastPrep-24 5G homogenizer (MP Biomedicals, Irvine, CA, USA). Olympus BX53; 40X.

**Additional file 6**. Virus-specific primers designed based on the HTS data.

**Additional file 7**. Viral contigs, sequences, BLAST hits, coverage, and descriptions.

**Additional file 8**. Identification of Acyrthosiphon pisum and Frankiniella occidentalis sequencing reads in alfalfa pollen samples.

## References

1. Bennet CW. Seed transmission of plant viruses. Adv Virus Res. 1969;14:221–61.

2. Johansen E. Seed transmission of viruses: current perspectives. Annu. Rev. Phytopathol. 1994. 32:363–86.

3. Pagán I. Transmission through seeds: The unknown life of plant viruses. PLoS Pathog. 2022; 18(8):e1010707.

4. Card SD, Pearson MN, and Clover GRG. Plant pathogens transmitted by pollen. Australasian Plant Pathol. 2007; 36:455–461.

5. Bradamante G, Scheid OM, Incarbone M. Under siege: virus control in plant meristems and progeny. Plant Cell 2021; 33:2523–2537.

6. Collavino A, Medina R, Nome C, Zanini A, Di Feo L. Evidence of cassava common mosaic virus in cassava shoot apical meristems. Plant Pathol. 2023;72:883–997.

7. Bayr D, Plaza MP, Gilles S, Kolek F, Leier-Wirtz V et al. Pollen long-distance transport associated with symptoms in pollen allergics on the German Alps: An old story with a new ending? Sci. Total Environ. 2023; 881:163310.

8. Suchan T, Talavera G, Saez L, Ronikier M, Vila R. Pollen metabarcoding as a tool for tracking long-distance insect migrations. Mol. Ecol. Res. 2018; 19:149–162.

9. Tayal M, Bonilla FR, Powell G, Cieniewicz E. Pollen-borne ilarviruses of peach: biology, ecology, and disease management. J. Integ. Pest Manag. 2024; 15(1):23.

10. Mahillon M, Brodard J, Schoen R, Botermans M, Dubuis N, Groux R, et al. Revisiting a pollen-transmitted ilarvirus previously associated with angular mosaic of grapevine. Virus Res. 2024; 344:199362.

11. Frosheiser FI. Alfalfa mosaic virus transmission to seed through alfalfa gametes and longevity in alfalfa seed. Phytopathol. 1974: 64:102–105.

12. Pesik Z., Hiruki C, Chen MH. Detection of Viral Antigen by Immunogold Cytochemistry in Ovules, Pollen, and Anthers of Alfalfa Infected with Alfalfa Mosaic Virus. Phytopathology 78:1027–1032. 1988.

13. Nemchinov LG, Irish BM, Grinstead S, and Olga A. Postnikova OA. Characterization of the seed virome of alfalfa (Medicago sativa L). Virol J. 2023a; 20(1):96.

14. Nemchinov LG, Irish BM, Grinstead S, Postnikova OA. Alfalfa transcriptomic responses to the field pathobiome. Plant Biol. (Stuttg). 2025; 27:492–503.

15. Nemchinov L.G., Postnikova O.A., Wintermantel W.M., Palumbo J.C., Grinstead S. 2023b Alfalfa vein mottling virus, a novel potyvirid infecting Medicago sativa L. Virol. J. 2023b; 20, 284.

16. Bolger AM, Lohse M, and Usadel B. Trimmomatic: a flexible trimmer for Illumina sequence data. Bioinformatics. 2014; 30: 2114–2120.

17. Bankevich A, Nurk S, Antipov D, Gurevich A, Dvorkin M, et al. SPAdes: A new genome assembly algorithm and its applications to single-cell sequencing. J Comput Biol. 2012; 19: 455–477.

18. Altschul SF, Gish W, Miller W, Myers EW, Lipman DJ. Basic local alignment search tool. J Mol Biol. 1990; 215:403–10.

19. Bushnell, B. BBMap: A fast, accurate, splice-aware aligner. 9th Annual Genomics of Energy & Environment Meeting, Walnut Creek, CA, USA. 2014. https://www.osti.gov/servlets/purl/1241166.

20. Wood DE, Lu J, Langmead B. Improved metagenomic analysis with Kraken 2. Genome Biol. 2019; 20(1):257.

21. Nemchinov LG, Grinstead S. Identification of a novel isolate of alfalfa virus S from China suggests a possible role of seed contamination in the distribution of the virus. Plant Dis. 2020; 104:3115–3117.

22. Larsen RC. Diseases caused by viruses. In: Compendium of alfalfa diseases and pests, third edition. Eds: Samac DA, Rhodes LH, Lamp WO. The American Phytopathological Society. 2015; p. 66–71.

23. Boutanaev AM, Nemchinov LG. Genome-wide identification of endogenous viral sequences in alfalfa (Medicago sativa L.). Virol J. 2021;18(1):185.

24. Chiba, S, Kondo H, Tani A, Saisho D, Sakamoto W. et al. Widespread endogenization of genome sequences of non-retroviral RNA viruses into plant genomes. PLoS Pathog. 2011; 7: e1002146.

25. Dietzgen RG, Kondo H, Goodin MM, Kurath G, Vasilakis N. The family Rhabdoviridae: mono- and bipartite negative-sense RNA viruses with diverse genome organization and common evolutionary origins. Virus Res. 2016; 227:158–170.

26. Al-Shahwan IM, Farooq T, Al-Saleh MA, Abdalla OA, and Amer MA. First Report of Red clover vein mosaic virus Infecting Alfalfa in Saudi Arabia. Plant Dis. 2016; 100:539.

27. Al-Shahwan IM, Abdalla OA, Al-Saleh MA, Amer MA. Detection of new viruses in alfalfa, weeds and cultivated plants growing adjacent to alfalfa fields in Saudi Arabia. Saudi Journal of Biological Sciences. 2017; 24:1336–1343.

28. Natsuaki T, Natsuaki KT, Okuda S, Teranaka M, Milne RG, Boccardo G. et al. Relationships between the cryptic and temperate viruses of alfalfa, beet and white clover. Intervirol. 1986: 25, 69–75.

29. Xie, WS, Antoniw, JF, White, RF, Jolliffe. TH. Effects of beet cryptic virus infection on sugar beet in field trials. Annals of Applied Biol. 1994;124: 451–459.

30. Jo Y, Choi H, Lee JH, Moh SH, Cho WK. Viromes of 15 pepper (Capsicum annuum L.) cultivars. Int J Mol Sci. 2022; 23(18):10507.

31. Dahan J, Wolf YI, Orellana GE, Wenninger EJ, Koonin EV, Karasev AV. A novel flavi-like virus in alfalfa (Medicago sativa L.) crops along the Snake River Valley. Viruses. 2022; 14:1320.

32. Hull R. Matthews’ Plant Virology. Fourth edition, Academic Press. 2002.

33. Harth JE, Winsor JA, Weakland DR, Nowak KJ, Ferrari MJ, Stephenson AJ. Effects of virus infection on pollen production and pollen performance: Implications for the spread of resistance alleles. Am J Bot. 2016;103:577–83.

34. Rajasekharan, PE, Kumar NKK, Harsha R, Mahapatra S, Vishwakarma PL. Effect of viruses infection symptoms on pollen viability in different horticultural crop species. Discover Plants. 2024; 1:18.

35. Pesic Z and Hiruki C. Effect of alfalfa mosaic virus on germination and tube growth of alfalfa pollen. Can. J. Plant Pathol. 1988; 10:6–10.

