## Additional files 1,2,3,4,5,6,8 for "Characterization of the alfalfa pollen virome"

**Additional file 1**

Alfalfa (*Medicago sativa* L.) plant genotype information used for sourcing pollen from field-grown plants in Prosser, WA.

| No. | Accession[1] | Name | Origin | Improvement level |
|---|---|---|---|---|
| 1 | PI 517233 | ILCA 5674 | Syria | Wild material |
| 2 | PI 552545 | UC 222 | United States | Breeding material |
| 3 | PI 516870 | Demnate | Morocco | Landrace |
| 4 | PI 536526 | MNGRN-2 | United States | Cultivar |
| 5 | PI 251560 | No. 1 | Former Serbia & Montenegro | Cultivated material |
| 6 | PI 516796 | GR 643 | Morocco | Landrace |
| 7 | PI 233199 | G 13621 | Russian Federation | Uncertain status |
| 8 | PI 399535 | Hunter River | Australia | Cultivar |
| 9 | PI 632163 | P.F. 5866 | Chile | Cultivated material |
| 10 | PI 231765 | G 5184 | United States | Uncertain status |
| 11 | PI 251693 | No. 22503 | Russian Federation | Uncertain status |
| 12 | PI 231042 | No. 4 | India | Uncertain status |
| 13 | PI 320535 | Crioula | Brazil | Landrace |
| 14 | PI 233195 | G 13620 | Russian Federation | Uncertain status |
| 15 | PI 255962 | Rambler | Canada | Cultivar |

[1] Individual plants (genotypes) were selected in prebreeding efforts from the plant introduction (PI) accessions which were originally obtained from the USDA ARS National Plant Germplasm System (NPGS).

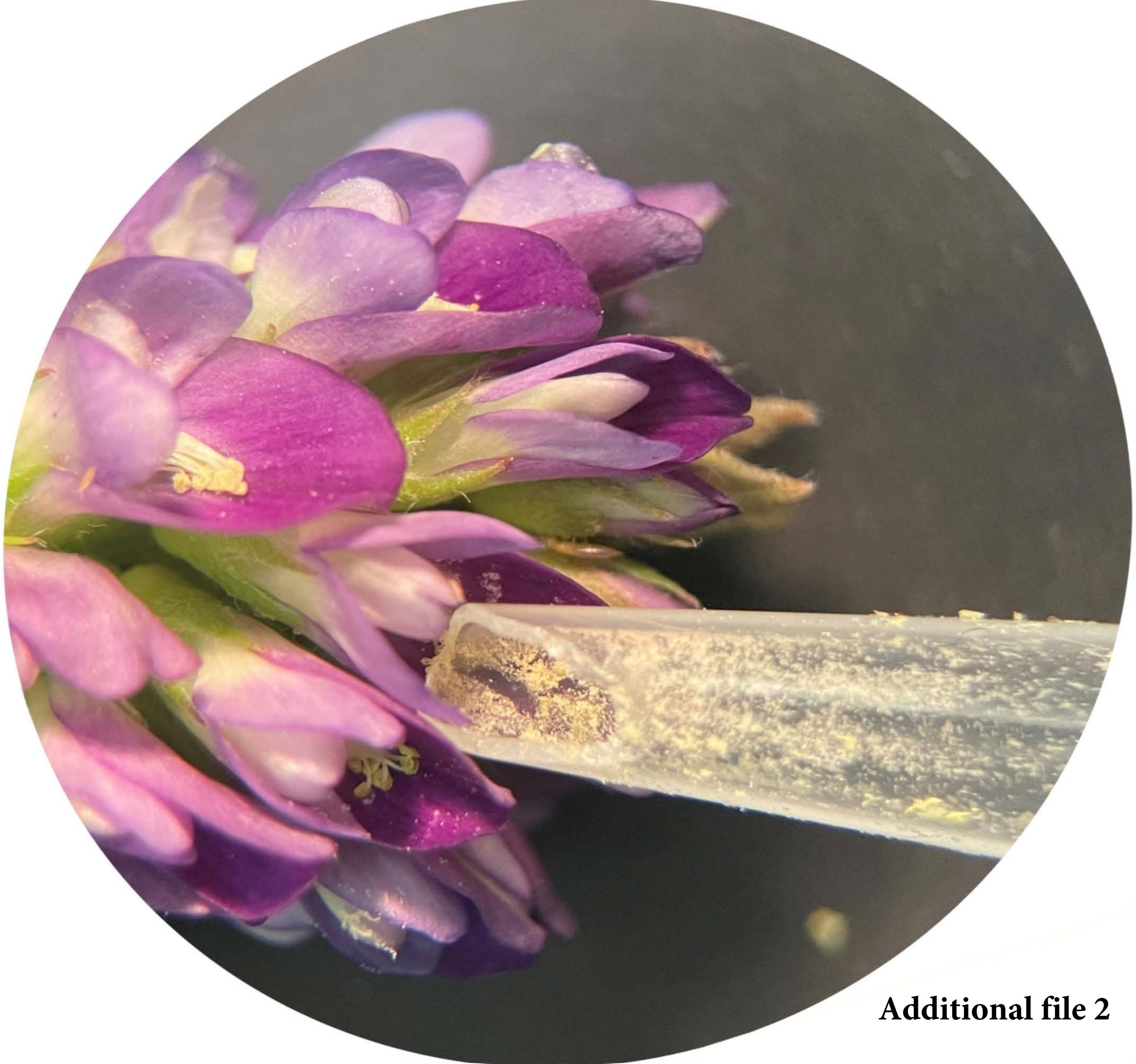

**Additional file 2**

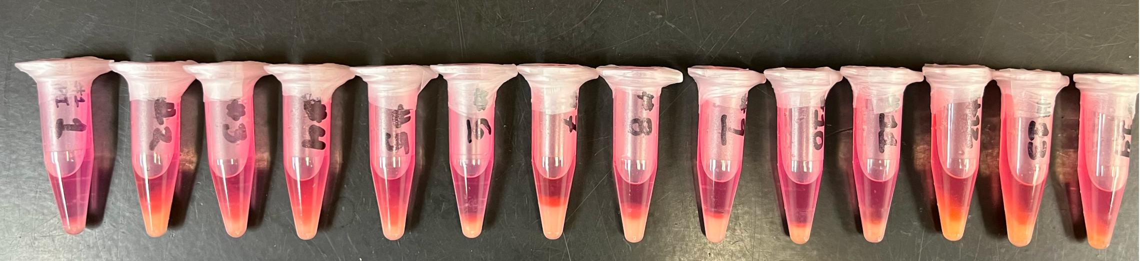

Additional file 3

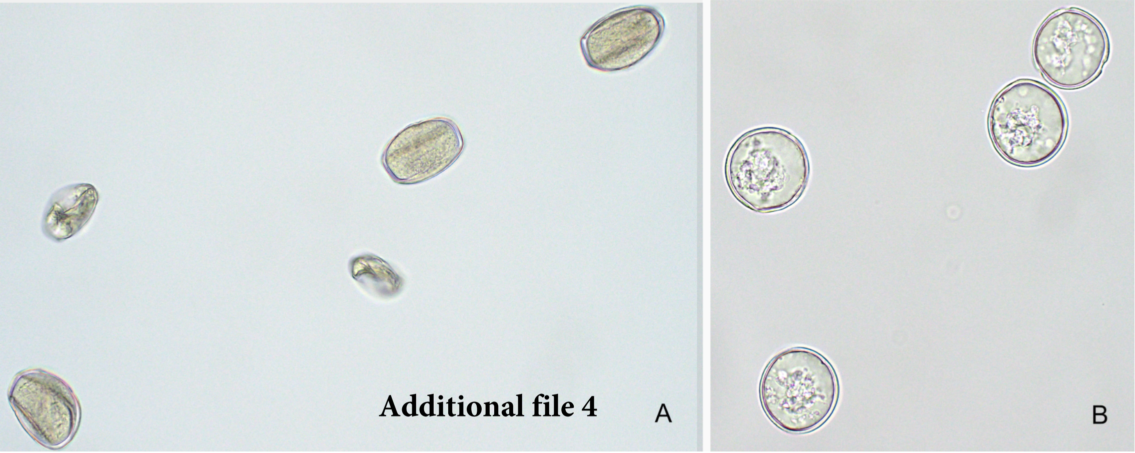
Additional file 4
A
B

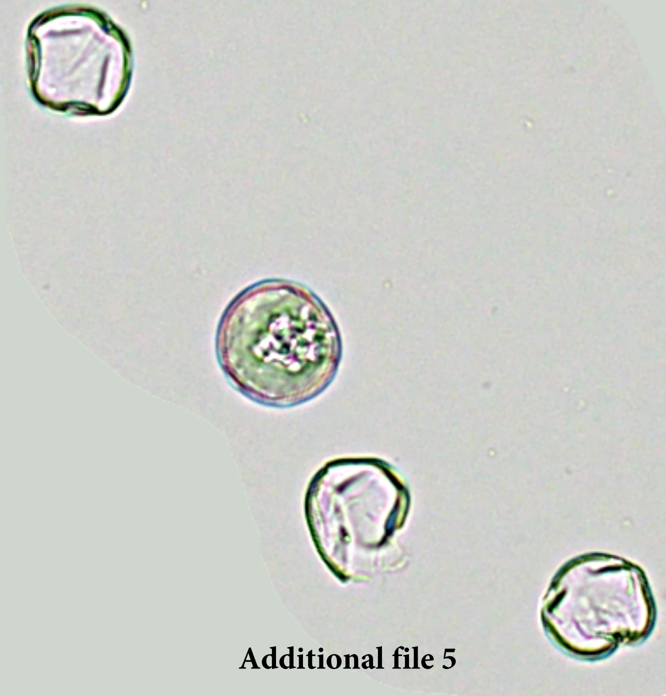

**Additional file 5**

**Additional file 6**

Virus-specific primers designed based on the HTS data

| Name | Sequence | Product length |
|---|---|---|
| PAPLV-F | TGGGAGAAGTAGAAGAGGAAGTA | |
| PAPLV-R | CCAGGCACACCTGATGTATAA | 314 |
| RCVMC-F | TGCTGTAGTAAGTGTGGTAAGTG | |
| RCVMC-R | GGTCATACCTTAAGCACCAAGA | 210 |
| BCV-F | CGTAGCCGTAAGAGGCTTG | |
| BCV-R | AGAACATTGGTGAAAGTGAGGA | 207 |
| BLRV-F | GAGATGCTATTGTTAAAGATGTTGGATG | |
| BLRV-R | AGAACATTGGTGAAAGTGAGGA | 205 |
| PeSV-F | GCCTCACGTTGGAGACAATA | |
| PeSV-R | CCCTCAGCATCCCGAATAAA | 554 |
| AVS-F | GGCCTTTACCAAGACGGTAATA | |
| AVS-R | GTAGCGGTGATTGTTGGATTTG | 639 |
| SRAV-F | GAGTCGTTGGCTAAGGTGATAC | |
| SRAV-R | GCCGCTAAACCTACTCCTATTC | 407 |
| ANRV-F | CCCACCTAAGCCACCTTTAAT | |
| ANRV-R | CAGAGCTACAGAGGGTGATTTC | 576 |
| MsAV1-F | AGTGTGGAGGACCCATTTATTC | |
| MsAV1-R | GGGTGGTGATGTAGCCAATTA | 317 |
| MsAPV1-F | GGATGAACTCGACCCTAAGAAC | |
| MsAPV1-R | CAAGCCCGACGAAAGTAGAA | 303 |

**Additional file 8**

Identification of *Acyrthosiphon pisum* and *Frankiniella occidentalis* sequencing reads in alfalfa pollen samples.

| Samples | [1]*A. pisum* reads | [2]*F. occidentalis* reads | Total sample reads |
|---|---|---|---|
| LN01 | 75 | 16131 | 81613114 |
| LN02 | 90 | 1844 | 75906416 |
| LN03 | 55 | 1330 | 86806392 |
| LN04 | 50 | 6172 | 66130554 |
| LN05 | 282 | 1322 | 82527766 |
| LN06 | 80 | 175 | 68517156 |
| LN07 | 36 | 738 | 63695260 |
| LN08 | 86 | 9881 | 72679330 |
| LN09 | 97 | 195 | 83426152 |
| LN10 | 103 | 12345 | 72358236 |
| LN11 | 218 | 27929 | 88360804 |
| LN12 | 93 | 351 | 100416738 |
| LN13 | 42 | 5911 | 56290254 |
| LN14 | 129 | 29934 | 57653144 |
| LN15 | 85 | 10743 | 71337634 |
| Average | 101.4 | 8333.4 | 75181263.33 |
| Part/total | 1.34874E-06 | 0.000110844 | |
| 1, *Acyrthosiphon pisum* | | | |
| 2, *Frankliniella occidentalis* | | | |
